# An engineered platform to study the influence of extracellular matrix nanotopography on cell ultrastructure

**DOI:** 10.1101/2025.02.27.640302

**Authors:** Shani Tcherner Elad, Rita Vilensky, Noa Ben Asher, Nataliya Logvina, Ran Zalk, Eyal Zussman, Leeya Engel

## Abstract

Nanoscale fabrication techniques have played an essential role in revealing the impact of extracellular matrix (ECM) nanotopography on cellular behavior. However, the mechanisms by which nanotopographical cues from the ECM influence cellular function remain unclear. To approach these questions, we have engineered a novel class of nanopatterned ECM constructs suitable for cryogenic electron tomography (cryo-ET), the highest resolution modality for imaging frozen hydrated cells in 3D. We electrospun aligned and randomly oriented ECM fibers directly onto transmission electron microscopy (TEM) supports to generate fibrous scaffolds that mimic physiological ECM in healthy (organized ECM) and diseased (disorganized ECM) states. We produced fibers from gelatin without toxic additives and crosslinked them to maintain structural stability in aqueous environments. The electrospun fibers had an average fiber diameter of hundreds of nanometers, within the physiological range. We confirmed that the TEM supports can serve as viable cell culture substrates that can influence cell organization and demonstrated their compatibility with plunge freezing and cryo-ET. By enabling nanoscale structural analysis inside cells on substrates with programmable topographies, this platform can be used to study the physical cues necessary for healthy endothelial tissue formation and pathologies that are linked to endothelial dysfunction in diseases such as peripheral arterial disease.

## 1 Introduction

Engineered ECM microenvironments have dramatically advanced our understanding of how micro- and nano-scale features guide cell organization and modulate cellular function ^1,2^. Nanotopographical cues influence cell polarity, adhesion ^3^, migration ^4^, proliferation ^5^, and differentiation ^6^. Disruptions to ECM structure can negatively impact development, tissue repair, and disease progression ^7^. In endothelial cells, which line all blood and lymphatic vessels, aligned nanofibrillar scaffolds have been shown to induce parallel alignment to mimic the elongated, atherosclerosis-resistant phenotype of cells within straight vessel segments ^8,9^. This nanofibril alignment reorganizes the cytoskeleton along the nanofibril direction and results in an endothelial phenotype that is resistant to inflammation and flow-induced shear stress ^10^.

While micro- and nanostructured cell culture scaffolds inform tissue engineering design choices, there remain gaps in our fundamental understanding of the mechanisms by which nanotopographical cues from the ECM modulate cell behavior and organization ^11,12^. Combining high-resolution imaging techniques with engineered ECM constructs can contribute to closing this knowledge gap.

Cryo-ET is a rapidly developing cryogenic transmission electron microscopy (TEM) technique capable of visualizing the interior of intact cells in 3D at the nanoscale without the need for fixation, dehydration, or staining ^13–16^. This technique is revolutionizing our structural understanding of the cell interior with major implications for mechanobiology, for example in elucidating the architecture of the cytoskeleton ^17^ and cell-ECM contacts ^18,19^.

The substrates that are used to culture cells for cryo-ET, termed EM grids, were developed for cryo-TEM of purified proteins and are coated with synthetic porous thin films that offer none of the rich physical cues that are present in vivo ^20,21^. We and others have developed methods to micropattern commercially available EM grids with two-dimensional islands of ECM proteins to direct cell shape and position cells for structural analysis with cryo-ET ^22–25^. EM grid micropatterning has facilitated cryo-ET of endothelial cells ^23^, however it relies on photo-patterning and is thus limited to a resolution of approximately 1–1.5 *µ*m. In addition, while ECM micropatterning can recapitulate certain mechanical aspects of the cell microenvironment (e.g., spatial constraints from neighboring cells), it cannot modulate substrate nanotopography.

Here we present a new class of nanofibrillar supports customized for imaging frozen hydrated cells with cryo-ET (see Fig. 1). We generate nanofibers using electrospinning and align them directly onto EM grids with a rotating collection disk ^26^. Our ability to tune the degree of organization of the gelatin nanofibers by varying the rotation speed of the disk faciliates 3D high-resolution imaging of the interior of endothelial cells cultured on thin films with different nanotopographies. This platform can be utilized to study the nanoscale underpinnings of cellular sensitivity to substrate nanotopography as well as the downstream effects of disruptions to ECM organization on cell ultrastructure.

**Fig. 1.**
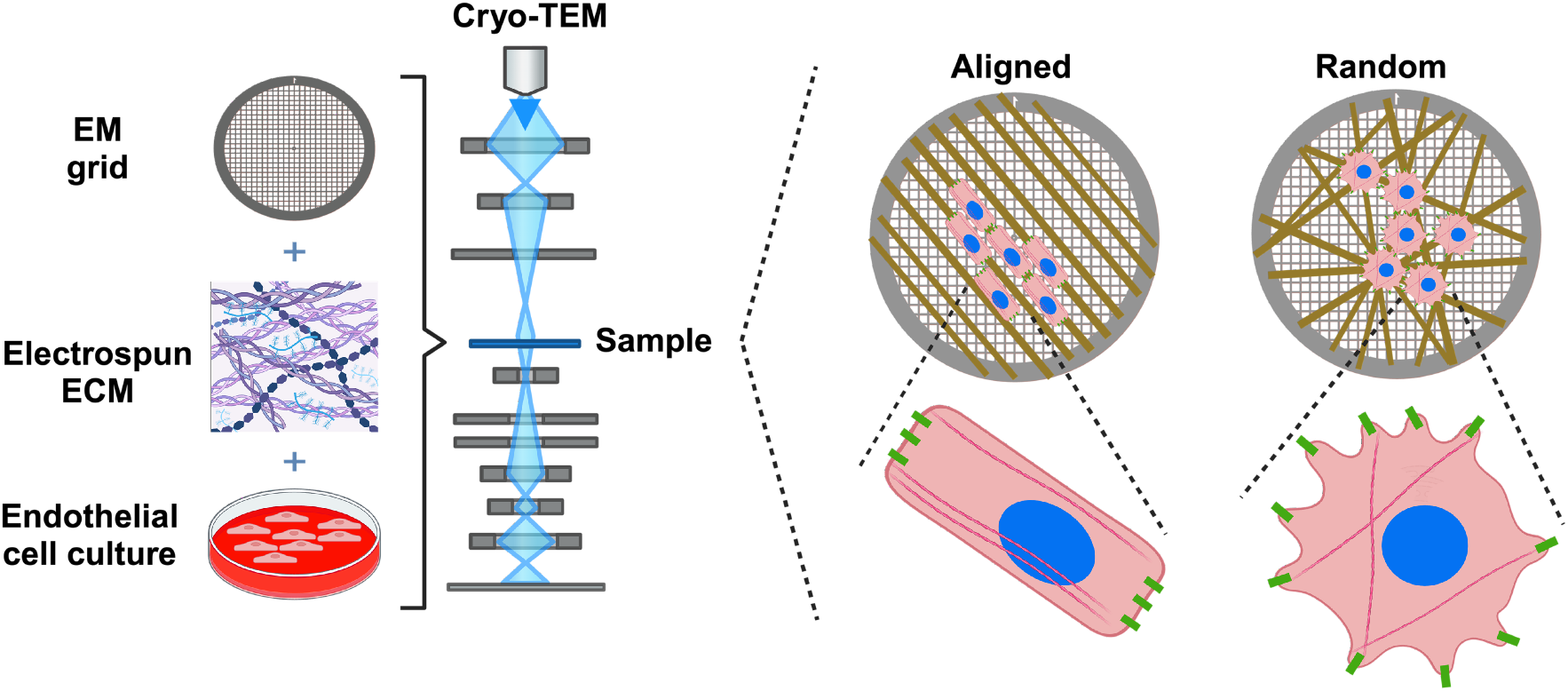
Schematic depicts a platform for high resolution cryo-TEM imaging of cells grown on ECM with different topographical cues.

## 2 Materials and Methods

### 2.1 Electrospinning gelating on EM grids

We electrospun gelatin fibers on gold grids coated with a porous gold thin film (UltrAuFoil R2/2, Quantifoil). EM grids are available in a range of conductive materials such as copper, but gold is preferred for cell culture due to its low toxicity. Grids with gold foil are also known to decrease the movement of frozen specimens during imaging relative to non-conductive foils such as carbon ^27^. We cleaned the grids using atmospheric plasma treatment for 30 seconds, assisted by a custom holder (see Fig. 2). To secure the grids during plasma exposure, we applied polydimethylsiloxane (PDMS) in a 1:10 ratio on top of a scanning electron microscopy (SEM) stub and cured it at room temperature for 24 hours.

**Fig. 2.**
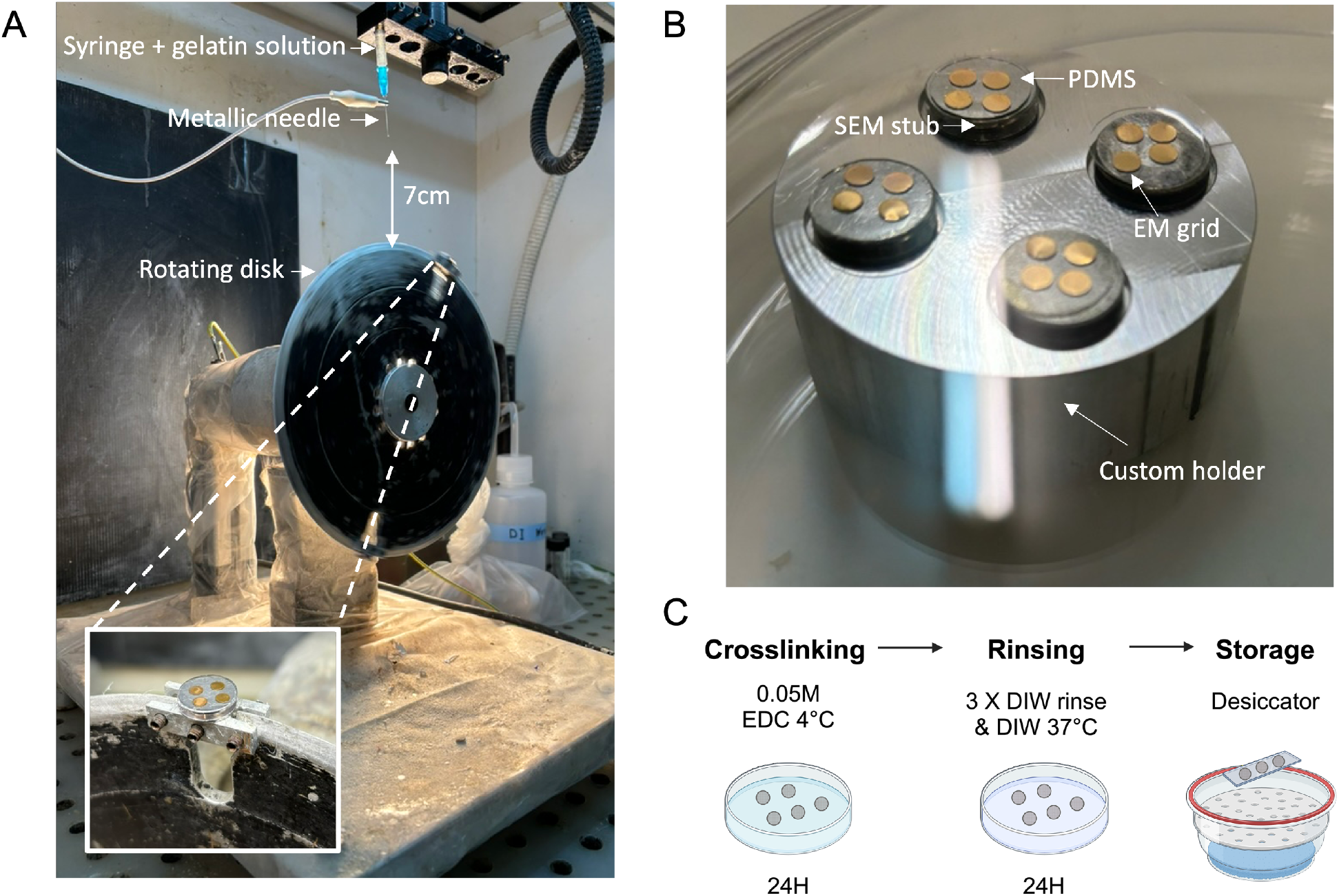
Experimental setup for depositing gelatin fibers on EM grids. (A) The electrospinning setup. Gelatin is extruded through a syringe needle under an electric field between the tip and the metal collector disk. The metal collector disk holds two SEM stubs coated with PDMS holding four EM grids in place (closeup in inset) (B). For plasma coating prior to electrospinning, the grids on the stubs are held upright using a custom aluminum holder. (C) EM grids coated with fibers are crosslinked, rinsed in water, and stored in a desiccator.

Electrospinning is a versatile method for producing polymer fibers by subjecting droplets of viscoelastic polymer solution to high voltage electrostatic fields ^28^. It has been applied to generate fibrous scaffolds for tissue engineering ^18,29,30^. Electrospun fibers can exhibit beads whose formation depends on the viscoelasticity of the solution, the charge density carried by the jet, and the surface tension of the solution ^31^.

We produced smooth fibers from a 28% gelatin solution made of 0.84 g of porcine skin-derived gelatin (Sigma-Aldrich, Type A, Bloom Strength 170-190 g) dissolved in 0.6 mL of acetic acid and 2.4 mL of DI water (8:2 v/v) ^32,33^. The solution was stirred for 30–60 minutes on a hot plate maintained at approximately 39 °C until electrospinning. It is essential to keep the solution warm to prevent gelation. Initially, a 30% gelatin solution was used in our experiments, however, the solution was excessively viscous and difficult to spin. Reducing the gelatin concentration to 28% yielded a stable electrospinning process with a smooth fiber morphology and free of “beads-on-string” type defects ^34^.

Electrospinning was performed using a rotating collector made of an aluminum disc with a diameter of 20 cm. The sharp edge of the disc serves to focus the electrical field and direct the polymer jet onto the EM grid that is affixed to it (see setup in Fig. 2) ^26^. Typically, two stubs holding eight grids were used per experiment. A 1 mL syringe with a 21 g needle (0.337 mm inner diameter) was employed for gelatin extrusion. The needle-to-grid distance was maintained at 7 cm. The potential difference between the syringe tip and the wheel was maintained at 13 kV with a flow rate set to 0.3 mL/hr. For the collection of aligned fibers, the tangential velocity of the disk was 11 m/s for a duration of 3.5 minutes; for randomly oriented fibers, it was set to 2.2 m/s for 2.5 minutes.

Experiments were conducted at room temperature (23±1 °C) and relative humidity of 67±4%. Following electrospinning, a heated metal biopsy punch (approximately 4 mm in diameter) was used to carefully excise the gelatin fibers around each grid, facilitating grid removal without damaging the fibers.

### 2.2 Crosslinking

To prevent the gelatin fibers from dissolving in an aqueous environment they must be crosslinked (see Fig. S1). Prior to electrospinning, a 0.05 M solution of N-(3-Dimethylaminopropyl)-N-ethylcarbodiimide hydrochloride (EDC) was prepared by dissolving 0.2 g of EDC in 16.7 mL of ethanol and 4.17 mL of deionized (DI) water (8:2 v/v). The solution was then stored at 4 °C. After electrospinning, the grids were immersed in the cold EDC 0.05 M solution and left overnight at 4 °C in glass dishes sealed with Parafilm ^35^. It is critical that the EDC solution remain cold during the crosslinking process ^35^.

To remove crosslinking residues, we created an array of three DI water drops (100–200 *µ*L each) on a Parafilm sheet. Grids were gently passed through the drops before being transferred to a glass Petri dish filled with DI water and incubated at 37 °C for 24 hours for the final rinse step ^35^.

### 2.3 Characterization of gelatin fibers

We utilized high-resolution scanning electron microscopy (HRSEM) to analyze gelatin fiber morphology and topographic structure (Zeiss Ultra Plus HRSEM). First we deposited an ultrathin layer of conductive metal onto the specimens by sputter-coating to prevent charging during imaging. We chose iridium for sputter coating, due to its high secondary electron yield and resistance to oxidation. The sputtered film thickness ranges from 5 to 7 nm, sufficient for excellent conductivity without significantly altering fiber topography.

We used Fiji software ^36^, along with the Directionality plugin, to analyze fiber orientation ^37^. This plugin finds the preferred fiber orientation by analyzing the input image and computing a directional histogram highlighting structures’ predominant orientation. The peaks indicate the preferred direction. Fiber diameters were measured manually using Fiji software.

### 2.4 Cell culture

Human umbilical vein endothelial cells (HUVECs) were cultured in EGM-2 MV Microvascular Endothelial Cell Growth Medium, supplemented with penicillin, streptomycin, and various growth factors. Cells were grown on dishes precoated with 0.2% gelatin and were maintained at 37 °C and 5% CO_2_. Experiments were carried out using cells from passages 7 to 12^23^.

Before cell seeding, we prepared glass-bottom dishes with a custom silicone sticker, creating wells that were 4 mm in diameter. We placed the grids fiber-coated side up in 20–30 *µ*L droplets of medium in the wells. We then added 5 µL from a cell suspension of approximately 1 million cells/mL, with the aim of depositing approximately two cells per grid-square. After the cells began to adhere to the fibers, 2 mL of warm media was added to each dish and they were left in the incubator overnight at 37 °C and 5% CO_2_.

### 2.5 Immunofluorescence

To fix the cells, we prepared a 4% paraformaldehyde (PFA) solution in PBS from a 16% PFA stock. After aspirating the culture media, the sample was rinsed with PBS to remove residual media. Following washing, we incubated the sample in 4% PFA at room temperature for 15 minutes, then rinsed it three times with PBS. The samples were then stored at 4 °C, in PBS, until immunostaining.

For immunostaining, cells were permeabilized using an Antibody Dilution Buffer (ADB) containing 1% bovine serum albumin (BSA) and 0.1% Triton X. Following a 1-hour blocking step with ADB, anti-paxillin staining (Abcam, ab32084, used at 1:200) was applied for three hours. After rinsing, a secondary staining solution containing anti-rabbit Alexa Fluor 488 conjugate antibody (Cell signaling 4412, used at 1:500) and ActinRed 555 (Thermo-Fisher Scientific R37112, 2 drops were used in 500 *µ*L of ADB) was applied for overnight incubation at 4 °C, followed by further rinses with PBS. Cells were imaged on an Olympus IX83 inverted microsope, using 20x and 40x objectives.

### 2.6 Vitrification, cryo-ET, and tomogram reconstruction

We vitrified the grids in a Leica EM GP2 plunge freezer, using a blotting time of 13 seconds. We dipped each grid in a droplet of PBS prior to loading it into the tool. Following freezing, we clipped the grids into autogrids and loaded them into a Thermo-Fisher Scientific Glacios cryogenic transmission electron microscope (cryo-TEM) with an X-FEG electron source operated at 200 kV. Tilt series were acquired in counting mode at a calibrated pixel size of 3.063 Å (39,000x nominal magnification) by a Falcon 4i direct detector coupled to a Selectris X energy filter set to ± 5 eV around zero-loss peak. Fiducials-free, dose-symmetric tilt series of 61 exposures from −60° to +60° at 2° intervals and a total dose of ∼120 ē /Å^2^ were collected with Tomo (Thermo-Fisher Scientific) at a defocus range between −2 *µ*m and −9 *µ*m. We reconstructed the tomograms using patch tracking in IMOD software package, version 4.9.1^38^. We segmented the tomogram volume and prepared the images in Fig. 5C using Amira (FEI/Thermo Fisher Scientific) software.

## 3 Results and Discussion

We successfully engineered nanopatterned ECM constructs that are compatible with cryo-ET by electrospinning 28% gelatin on gold EM grids (Fig. 3). Plasma treating the EM grids prior to electrospinning was critical for maintaining adhesion between the gelatin fibers and the gold EM grid surface. Crosslinking the gelatin fibers was critical to preventing their dissolution in (aqueous) cell culture medium (Fig. S1).

**Fig. 3.**
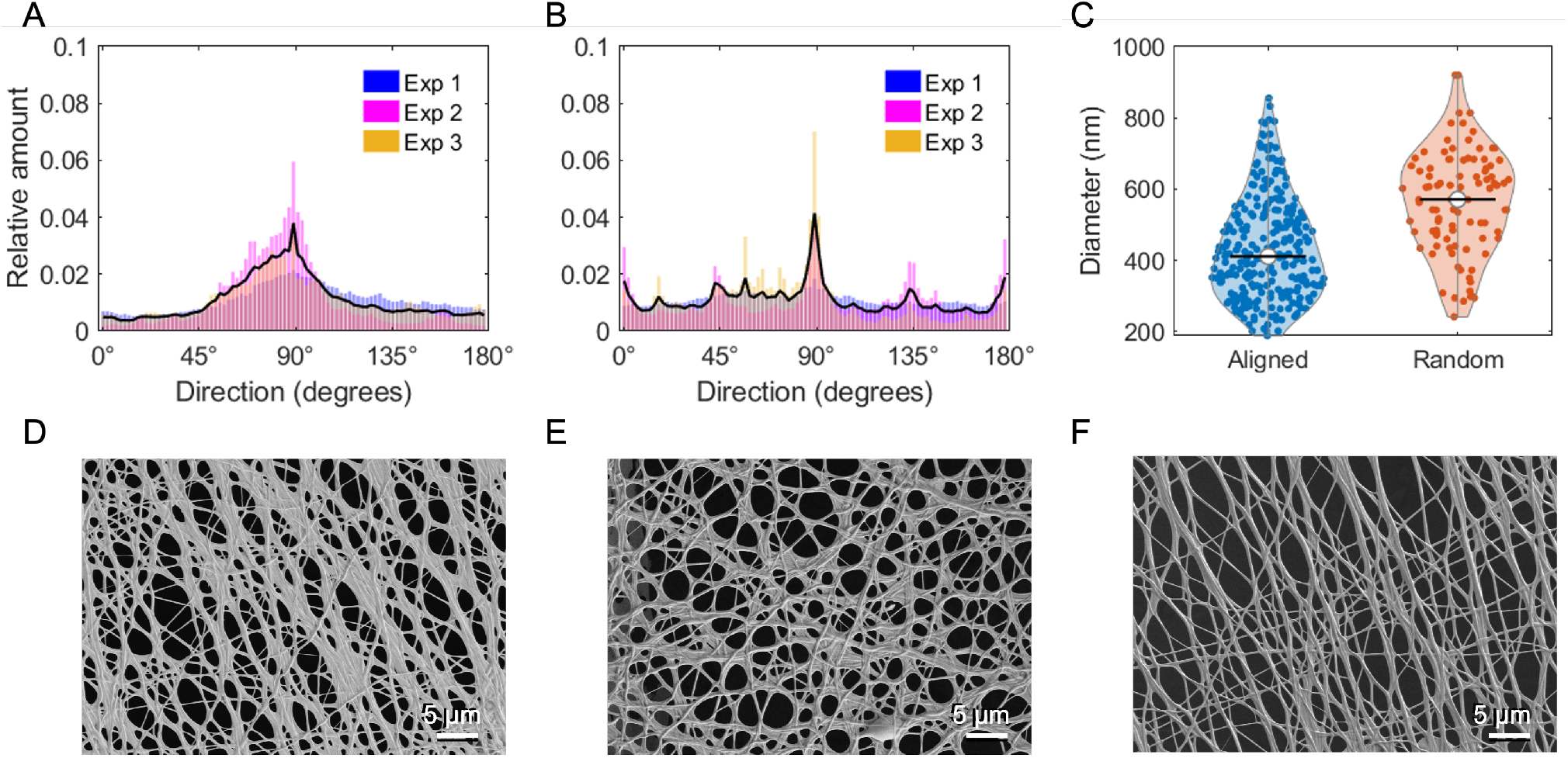
Fiber characterization. (A-B) Histograms of preferred fiber orientation for three separate experiments for aligned (A) and random (B) fiber orientations. The black line represents a weighted average for the three experiments for each condition. (C) Violin plot depicts fiber diameter for random (red) and aligned (blue) oriented fibers. The black line indicates the median. (D-F) HRSEM images of gelatin fibers electrospun from 28% solution directly onto gold TEM grids. (D) Aligned fibers. (E) Randomly oriented fibers. (F) Aligned oriented fibers after 4 months of storage in a desiccator.

We observed a higher degree of fiber orientation when the collection wheel was rotating at a faster rate (aligned condition) versus the slower rate of rotation (random condition) (Fig. 3A and B, respectively and Fig. S2). We measured an average fiber diameter of 417.7 +/-9.0 nm (SEM) for the aligned condition and 479.6 +/-10.3 nm (SEM) for the random condition (Fig. 3C).

In vivo, ECM is composed of various proteins that can bundle into larger secondary structures ranging from 200 nm to 1 *µ*m in diameter ^39^, thus the gelatin fibers we produced are within the physiological range. The slight disparity in diameter between the conditions may be due to the fibers in the aligned condition undergoing more elongation due to tensile stress as the rotating disk collected the fibers at a higher tangential velocity for that condition. EM grids with electrospun fibers had a shelf-life of at least four months when stored in a desiccator (Fig. 3F).

The electrospun EM grids serve as viable cell culture platforms. Our immunofluorescence results show that the shape and orientation of endothelial cells grown on the electrospun TEM grids are influenced by the fiber direction (Fig. 4), consistent with previous reports of cells cultured on engineered aligned nanofibrillar ECM constructs ^9,10^. On the aligned fibers, the cells display an elongated morphology and align with the fiber direction. In contrast, on the randomly oriented fibers, the cells adopt a rounder, star-shaped morphology. F-actin-rich filopodia appear to follow the orientation of the gelatin nanofibers.

**Fig. 4.**
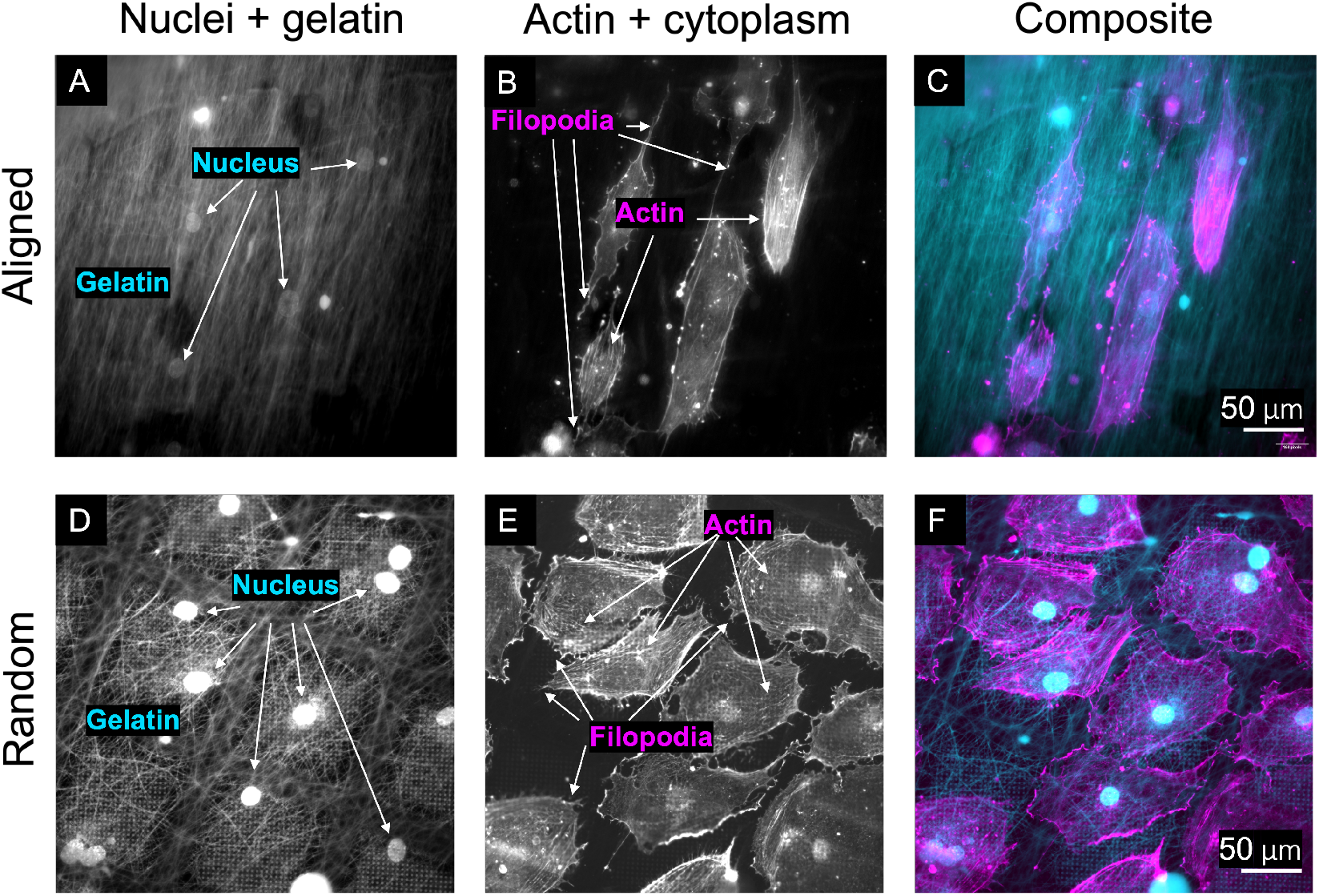
Immunofluorescence of endothelial cells grown on EM grids coated with aligned (A-C) and randomly (D-F) oriented gelatin fibers for 24 hours. RFP-expressing cells were stained for DNA and F-actin. The aligned (A) and randomly oriented (D) gelatin fibers are autofluorescent and thus apparent in the 405 nm channel with the nuclei. (B-E) F-actin, along with some background signal from the cytoplasm of the cells. (C-F) composite images of F-actin (magenta), nuclei, and gelatin (cyan) show that the orientation of the nanofibers influences the cell morphology and the orientation of F-actin-rich filopodia.

**Fig. 5.**
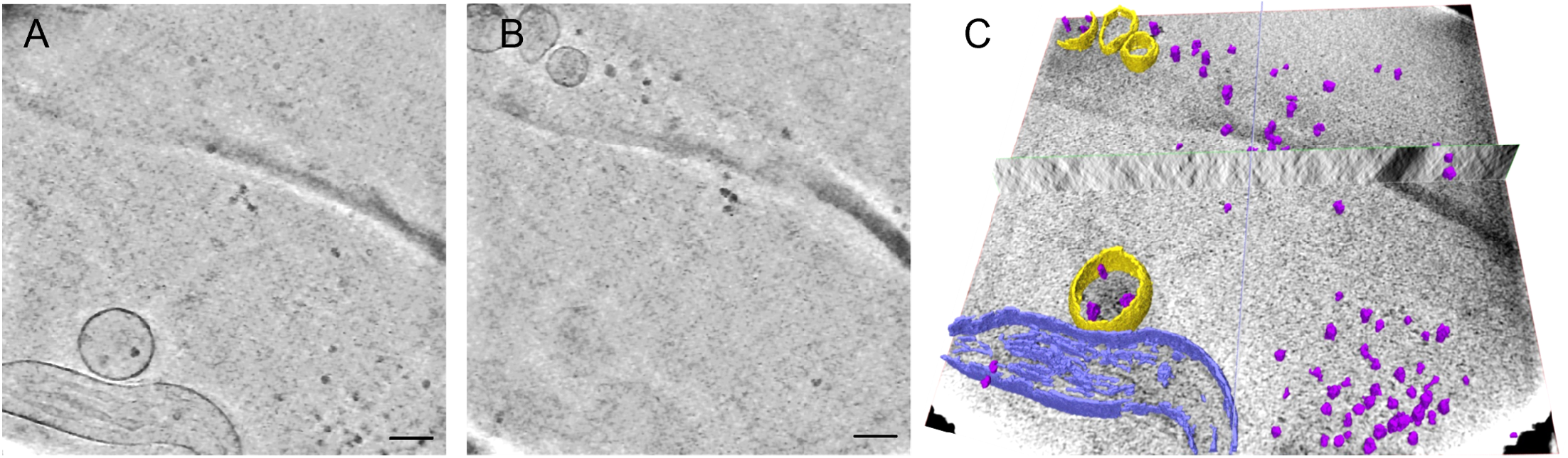
(A,B) Slices from a cryo-tomogram of endothelial cell cultured on electrospun gelatin fibers. Scale bars, 100 nm. (C) Segmented features overlaid with tomographic slice depicting mitochondria (blue), granules or ribosomes (violet), and vesicles (yellow).

We demonstrate that our engineered ECM constructs are compatible with vitrification by plunge freezing and cryo-ET (see Fig. 5 and Supplementary Movies 1-3). Despite the added gelatin, the grids were readily clipped into standard autogrids for automatic handling within the cryo-TEM where they were imaged. Our reconstructed tomograms show cellular features such as mitochondria, granules, ribosomes, and vesicles in 3D, some of which are segmented in Fig. 5 (see also Supplementary Movie 1). In this study we were limited to imaging thin (several 100s nm thick) regions of the cells, especially because we were using a 200 kV (vs. 300 kV) microscope ^40,41^. Cryo-focused ion beam milling (cryo-FIB) can be used in future studies to generate thin lamellae in the frozen cells to facilitate their imaging with cryo-ET ^42,43^.

Previous methods for producing aligned collagen fibrillar films, such as extrusion, are not readily adapted to cryo-TEM because resulting ECM mats are on the order of hundreds of micrometers thick. Electrospinning allows us to deposit a layer of nanofibers on the EM grids that is sufficient to orient the cells, but thin enough for vitrification by plunge freezing (<10 *µ*m).

This new class of cryo-ET cell culture supports can be used to investigate the organization of the cell-ECM adhesions that underlie the mechanosensitivity of endothelial cells to changes in ECM topography, ultimately improving our understanding of the mechanical cues required for the formation of healthy tissue and of the pathologies that result from defects in endothelial tissue organization associated with peripheral arterial disease.

Studying the nanoscale structures at the cell-ECM interface can also contribute toward answering long-held questions in tissue engineering such as why certain fibril dimensions are better than others at promoting cell survival in engineered tissues. We anticipate that the fibers will provide a valuable means for locating and identifying the cell-ECM adhesions.

Finally, by mimicking the physical cues present in physiological ECM, these EM grids create a more in vivo-like environment that can improve the translational relevance of cryo-ET studies of mammalial cell ultrastructure and support the growth of cell types that are challenging to culture (e.g., tissue-derived primary cells or stem cells) on EM grids.

## Conclusions

We have developed a new class of supports for cryo-ET that are coated with thin films of aligned ECM fibers to provide cells with programmed nanotopographical cues that mimic in vivo ECM topography. We controlled the degree of alignment between the fibers in the ECM thin films by varying the electrospinning parameters, specifically the speed of the rotating collector wheel. This platform can be used in future cryo-ET studies to probe how physical cues in the cell microenvironment influence junctional and cytoskeletal architecture.

## Author contributions

**Shani Tcherner Elad**: Conceptualization, Methodology, Software, Formal analysis, Investigation, Data curation, Writing - Original Draft, Visualization, Project administration. **Rita Vilen-sky**: Methodology, Investigation, Writing - Review & Editing. **Noa Ben Asher**: Methodology, Investigation, Writing - Review & Editing. **Nataliya Logvina**: Software, Formal analysis, Data curation, Writing - Review & Editing, Visualization. **Ran Zalk**: Software, Investigation. **Eyal Zussman**: Resources, Writing - Review & Editing, Supervision. **Leeya Engel**: Conceptualization, Methodology, Investigation, Writing - Original Draft, Supervision, Project administration, Funding acquisition.

## Data availability

The data supporting this article have been included as part of the Supplementary Information. Tomograms and raw electron microscopy data have been uploaded to the EMPIAR database (accession number TBD).

## Acknowledgments

This work was supported by the Israel Science Foundation (ISF) Personal Research Grant no. 1925/23. We thank Dr. Ngan Huang and Dr. Doron Elad for helpful advice and Dr. Olga Kleinerman for HRSEM imaging, which was performed at the Technion Center for Electron Microscopy of Soft Matter. The authors are grateful for the generous support from the Guzik Foundation to BGU’s Cryoelectron microscopy unit, where cryo-ET was performed. L.E. is a Diane and Guilford Glazer Foundation Faculty Fellow. Figures 1 and 2C were created with Biorender.com.

