## Supplementary Information for "An engineered platform to study the influence of extracellular matrix nanotopography on cell ultrastructure"

### Supplemental Information

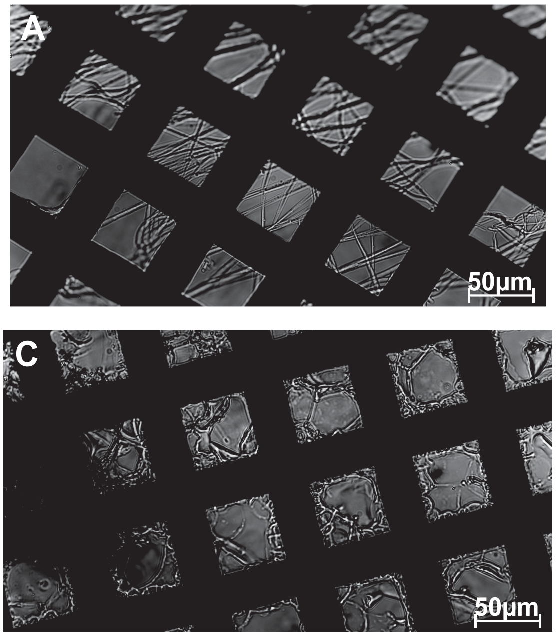

**Figure 1:** Micrographs of gelatin fibers after introduction to deionized water show that crosslinking (here with EDC) is critical for maintaining fiber structure. (A) Crosslinked 20% gelatin fibers with 50/50 v/v TFE/DI water. (B) Crosslinked 30% gelatin fibers with 2/8 v/v acetic acid/DI water. (C-D) Loss of structure without crosslinking. (C) 20% gelatin fibers with 50/50 v/v TFE/DI water. (D) 30% gelatin fibers with 2/8 v/v acetic acid/DI water.

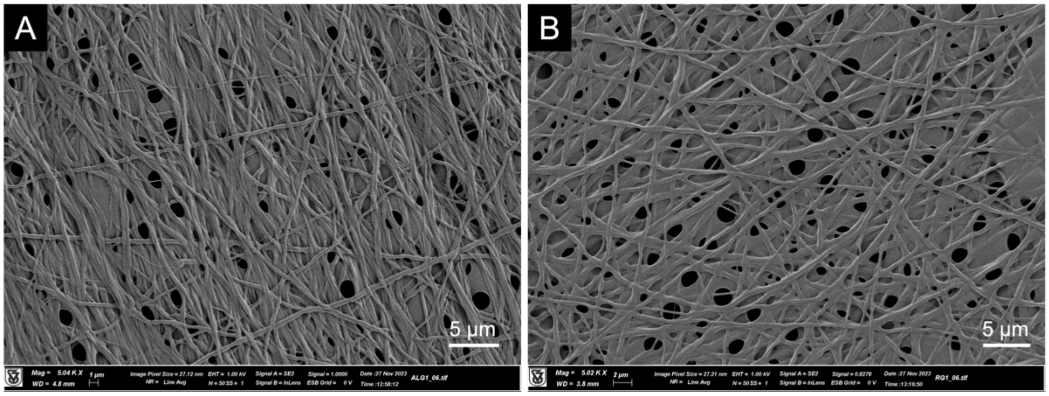

**Figure 2**: HR-SEM images of gelatin fibers electrospun from a 28% solution directly onto gold TEM grids (UltrAufoil 200 mesh R2,2). (A) Aligned fibers. (B) Randomly oriented fibers.

**Table 1:** Measured fiber diameters from HRSEM images.

| Aligned- exp1 | Aligned- exp2 | Aligned- exp3 | Random- exp1 | Ramdom- exp2 | Random- exp 3 |
|---|---|---|---|---|---|
| 649.434 | 294.365 | 394.447 | 464.145 | 505.139 | 459.61 |
| 613.031 | 328.2 | 406.992 | 792.829 | 658.221 | 481.857 |
| 626.93 | 484.334 | 278.487 | 352.804 | 814.277 | 484.334 |
| 446.266 | 294.365 | 341.777 | 467.741 | 439.783 | 484.581 |
| 652.743 | 440.327 | 218.8 | 559.119 | 601.998 | 542.386 |
| 626.548 | 356.181 | 280.201 | 536.395 | 683.903 | 610.487 |
| 402.259 | 356.181 | 402.259 | 437.6 | 479.866 | 470.547 |
| 455.293 | 364.156 | 333.266 | 855.841 | 434.306 | 562.32 |
| 293.551 | 333.266 | 358.859 | 235.655 | 541.723 | 418.732 |
| 570.561 | 285.28 | 364.156 | 201.724 | 920.261 | 380.548 |
| 249.47 | 364.156 | 201.724 | 109.4 | 738.106 | 240.929 |
| 493.635 | 526.94 | 442.496 | 140.1 | 681.449 | 285.28 |
| 716.716 | 313.274 | 264.377 | 473.337 | 510.325 | 188.854 |
| 443.036 | 345.953 | 401.663 | 323.054 | 650.539 | 497.259 |
| 294.365 | 503.715 | 503.715 | 206.416 | 634.52 | 571.085 |
| 503.715 | 360.854 | 303.967 | 372.603 | 616.535 | 761.883 |
| 717.717 | 382.118 | 418.017 | 437.6 | 461.559 | 313.465 |
| 543.488 | 560.828 | 403.447 | 394.447 | 704.59 | 350.08 |
| 788.895 | 309.43 | 176.402 | 465.175 | 636.027 | 339.316 |
| 631.495 | 341.777 | 451.068 | 499.899 | 616.535 | 535.166 |
| 568.88 | 491.692 | 188.219 | 402.259 | 785.55 | 372.603 |

| | | | | | |
|---|---|---|---|---|---|
| 503.715 | 493.635 | 323.054 | 536.395 | 715.379 | 507.503 |
| 328.2 | 480.364 | 313.274 | 387.715 | 626.548 | 308.073 |
| 574.324 | 540.396 | 352.804 | 507.503 | 541.281 | 606.85 |
| 401.663 | 424.833 | 498.941 | 796.444 | 686.697 | 430.569 |
| 232.588 | 249.47 | 609.506 | 559.119 | 714.374 | 517.081 |
| 266.182 | 267.079 | 342.476 | 629.977 | 610.291 | 372.121 |
| 649.803 | 325.27 | 364.156 | 712.361 | 418.017 | 476.989 |
| 784.33 | 278.487 | 451.068 | 244.626 | 570.561 | |
| 560.401 | 356.181 | 285.28 | 207.572 | 1103.802 | |
| 402.259 | 495.088 | 305.146 | 318.895 | 665.454 | |
| 529.207 | 430.986 | 240.68 | 557.731 | 605.171 | |
| 387.097 | 249.47 | 284.019 | 203.818 | 297.6 | |
| 461.559 | 232.588 | 502.049 | 561.591 | 294.365 | |
| 389.562 | 547 | 612.347 | 439.037 | 683.903 | |
| 754.464 | 424.833 | 411.96 | 472.927 | | |
| 341.777 | 477.866 | 332.727 | 478.516 | | |
| 574.324 | 464.145 | 356.181 | 404.772 | | |
| 376.438 | 280.201 | 449.739 | 455.75 | | |
| 541.281 | 263.47 | 441.005 | 694.178 | | |
| 553.957 | 280.201 | 280.414 | 561.591 | | |
| 728.64 | 244.626 | 445.461 | 641.168 | | |
| 529.659 | 418.017 | 300.999 | 628.605 | | |
| 389.562 | 479.866 | 337.725 | 490.519 | | |
| 394.447 | 285.28 | 674.121 | 506.846 | | |
| 520.082 | 541.281 | 397.62 | 495.91 | | |
| 643.51 | 529.659 | 388.486 | 168.072 | | |
| 485.815 | 294.365 | 604.082 | 543.853 | | |
| 418.017 | 218.8 | 411.233 | 531.491 | | |
| 516.387 | 232.588 | 483.593 | 486.437 | | |
| 622.716 | 465.175 | 323.979 | 461.547 | | |
| 833.166 | 403.447 | 363.662 | 575.618 | | |
| 371.96 | 451.068 | 264.829 | 473.278 | | |
| 540.396 | 267.079 | 263.47 | 415.708 | | |
| 667.967 | 652.743 | 406.108 | 490.858 | | |
| 722.371 | 185.658 | 270.862 | 452.824 | | |
| 307.1 | 201.724 | 308.849 | 555.641 | | |
| 479.866 | 325.27 | 318.014 | 541.403 | | |
| 550.924 | 267.079 | 365.632 | 518.194 | | |
| 754.464 | 681.449 | 349.053 | 439.037 | | |

| | | | |
|---|---|---|---|
| 416.43 | 293.551 | 418.017 | 412.831 |
| 330.35 | 256.099 | 457.13 | 291.916 |
| 568.909 | 855.841 | 505.139 | 510.794 |
| 525.184 | 185.658 | 519.736 | 746.81 |
| 709.788 | 236.668 | 485.568 | 339.669 |
| 750.731 | 333.266 | 389.562 | 387.097 |
| 502.64 | | 256.332 | 495.088 |
| 270.652 | | 161.527 | 658.221 |
| 590.189 | | 477.616 | 489.252 |
| 425.311 | | 531.125 | 403.447 |
| 347.981 | | 358.358 | 520.082 |
| 552.917 | | 296.189 | 614.201 |
| 794.743 | | 329.11 | 201.724 |
| 552.917 | | 429.038 | 297.6 |
| 411.01 | | 645.831 | 333.984 |
| 450.903 | | 717.884 | 280.201 |
| 599.613 | | 581.469 | 323.054 |
| 295.727 | | 416.583 | 399.273 |
| 330.35 | | 535.166 | 510.794 |
| 379.819 | | | 402.259 |
| 574.791 | | | 489.252 |
| 600.236 | | | 315.558 |
| 464.78 | | | 577.649 |
| 441.691 | | | 634.52 |
| 492.879 | | | 661.486 |
| 611.341 | | | 303.967 |
| 380.802 | | | 424.833 |
| 391.449 | | | 443.036 |
| 528.023 | | | 403.447 |
| 676.354 | | | 278.487 |
| 634.633 | | | 463.629 |
| 437.673 | | | 244.626 |
| 113.984 | | | 646.109 |
| 198.629 | | | 366.776 |
| 242.08 | | | 461.559 |
| 459.589 | | | 455.293 |
| 371.965 | | | 597.208 |
| 133.111 | | | 574.324 |
| 50.901 | | | 717.717 |

| | | | |
|---|---|---|---|
| 307.109 | | | 529.659 |
| 393.967 | | | 313.274 |
| 481.464 | | | 715.379 |
| 264.268 | | | 616.146 |
| 306.379 | | | 700.502 |
| 96.228 | | | 637.906 |
| 156.467 | | | 442.496 |
| 286.465 | | | 579.304 |
| 29.643 | | | 622.331 |
| 351.539 | | | 364.156 |
| 439.424 | | | 597.608 |
| 372.497 | | | 468.252 |
| 375.859 | | | 420.87 |
| 137.239 | | | 559.119 |
| 180.983 | | | 727.325 |
| 350.651 | | | 715.379 |
| 69.89 | | | 402.259 |
| 331.408 | | | 353.482 |
| 437.673 | | | 263.47 |
| 49.14 | | | 569.301 |
| 176.96 | | | 356.181 |
| 350.763 | | | 420.87 |
| 679.69 | | | 154.715 |
| 574.324 | | | 334.7 |
| 722.371 | | | 655.67 |
| 510.325 | | | 469.784 |
| 510.325 | | | 901.339 |
| 687.046 | | | 616.535 |
| 491.692 | | | 657.857 |
| 371.96 | | | 798.545 |
| 686.697 | | | 614.201 |
| 403.447 | | | 461.559 |
| 313.274 | | | 309.43 |
| 360.854 | | | 371.96 |
| 510.794 | | | 303.967 |
| 309.43 | | | 284.44 |
| 241.672 | | | 372.603 |
| 207.572 | | | 412.831 |
| 328.929 | | | 371.316 |

| | | | |
|---|---|---|---|
| 532.813 | | | 371.96 |
| 294.365 | | | 412.831 |
| 433.202 | | | 588.731 |
| 339.669 | | | 403.447 |
| 484.334 | | | 263.47 |
| 201.724 | | | 540.396 |
| 294.365 | | | 353.482 |
| 353.482 | | | 643.51 |
| 790.41 | | | 285.28 |
| 285.28 | | | 266.182 |
| 528.754 | | | 284.44 |
| 264.377 | | | 532.813 |
| | | | 416.295 |
| | | | 358.859 |
| | | | 437.6 |
| | | | 339.669 |
| | | | 315.558 |
| | | | 420.87 |
| | | | 704.59 |
| | | | 340.373 |
| | | | 402.259 |
| | | | 364.156 |
| | | | 577.649 |
| | | | 241.672 |
| | | | 249.47 |
| | | | 372.603 |
| | | | 159.289 |
| | | | 387.715 |
| | | | 442.496 |
| | | | 520.082 |
| | | | 636.027 |
| | | | 609.506 |
| | | | 455.293 |
| | | | 762.04 |
| | | | 350.08 |
| | | | 360.854 |
| | | | 422.573 |
| | | | 481.857 |
| | | | 225.268 |

| | | | |
|---|---|---|---|
| | | | 270.641 |
| | | | 389.562 |
| | | | 345.953 |
